# A performant bridge between fixed-size and variable-size seeding

**DOI:** 10.1101/825927

**Authors:** Arne Kutzner, Pok-Son Kim, Markus Schmidt

**Affiliations:** Department of Information Systems, College of Engineering, Hanyang University, 222 Wangsimni-ro, Seongdong-gu, Seoul, 04763, Republic of Korea; Department of Mathematics, College of Science and Technology, Kookmin University, 77, Jeongneung-ro, Seongbuk-gu, Seoul, 02707, Republic of Korea

**Keywords:** high-throughput sequence alignment, minimizer, SMEM, FMD-Index, seed entropy

## Abstract

**Background:** Seeding is usually the initial step of high-throughput sequence aligners. Two popular seeding strategies are fixed-size seeding (*k*-mers, minimizers) and variable-size seeding (MEMs, SMEMs, maximal spanning seeds). The former strategy supports fast seed computation, while the latter one benefits from a high seed entropy. Algorithmic bridges between instances of both seeding strategies are of interest for combining their respective advantages.

**Results:** We introduce an efficient strategy for computing MEMs out of fixed-size seeds (*k*-mers or minimizers). In contrast to previously proposed extend-purge strategies, our merge-extend strategy prevents the creation and filtering of duplicate MEMs. Further, we describe techniques for extracting SMEMs or maximal spanning seeds out of MEMs. A comprehensive benchmarking shows the applicability, strengths, shortcomings and computational requirements of all discussed seeding techniques. Additionally, we report the effects of seed occurrence filters in the context of these techniques.

Aside from our novel algorithmic approaches, we analyze hierarchies within fixed-size and variable-size seeding along with a mapping between instances of both seeding strategies.

**Conclusion:** Benchmarking shows that our proposed merge-extend strategy for MEM computation outperforms previous extend-purge strategies in the context of PacBio reads. The observed superiority grows with increasing read size and read quality. Further, the presented filters for extracting SMEMs or maximal spanning seeds out of MEMs outperform FMD-index based extension techniques. All code used for benchmarking is available via GitHub at https://github.com/ITBE-Lab/seed-evaluation.

## 1 Background

Most high-throughput read aligners [1-5] perform the following three steps: seeding [6, 7], seed processing (e.g. chaining) [8, 9] and dynamic programming [10, 11]. A sequence used as target for alignments is called reference. Reads aligned against such a reference are called queries. There are two techniques for seed computation: fixed-size seeding [12] and variable-size seeding [13, 14]. Fixed-size seeding is usually done via *k*-mers or their space efficient variant, *w, k*-minimizers, where *k*-mer seeds are perfect matches of size *k* between query and reference. Given a window size *w*, the set of *w, k*-minimizer seeds [3] is a space efficient subset of the set of *k*-mer seeds. Variable-size seeding relies on maximal exact matches (MEMs) or subsets of them. A MEM [6] is a perfect match between query and reference that cannot be extended further in either direction. MEMs can be computed directly via some form of full-text search index as e.g. the FM-index [13, 14] or the FMD-index [15]. There are two subgroups within MEMs: SMEMs (super-maximal exact matches) [15] and maximal spanning seeds [2]. A SMEM is a MEM that is not enclosed by another MEM on the query. A MEM is a maximal spanning seed if and only if it comprises at least one query position that it is not covered by another longer MEM. This implies that a maximal spanning seed is always a SMEM but not the contrary.

There exist proper subset relationships among variable-size seeding techniques as well as fixed-size seeding techniques. Further, there is a mapping between both groups as presented in Fig 1. Alg. 1 (Section 2.4) implements this mapping using a merge-extend strategy. The mapping is surjective, because for every MEM there is at least one corresponding *k*-mer seed. However, it is not injective, because several *k*-mer seeds can map to the same MEM. Trivially, the inverse mapping can be achieved by simply computing all *k*-size sub seeds of each MEM. *w, k*-minimizers are mapped to a subgroup within the set of MEMs.

**Figure 1.**
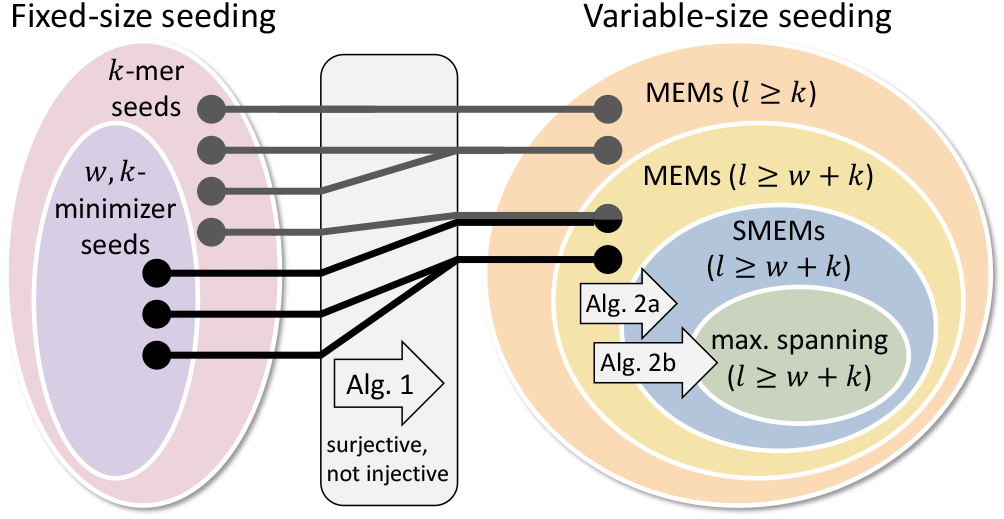
The hierarchies for fixed-size seeding (*k*-mer seeds and minimizer seeds) and variable-size seeding (MEMs, SMEMs, maximal spanning seeds) as well as the mapping among them. The ovals indicate subset relationships among the different seed sets. *l* denotes the length of an individual variable-size seed. Alg. 1 (Section 2.4) constructs MEMs of size ≥ *k* and ≥ *w* + *k* out of *k*-mer seeds and (*w, k*)-minimizer seeds, respectively. Alg. 2a (Section 2.6) in the methods section extracts SMEMs out of MEMs. Alg. 2b (Section 2.6) computes maximal spanning seeds from MEMs (or SMEMs).

An aligner can only find the correct location of a read if there are seeds for this location. Therefore, the seeding defines an upper bound for the maximal accuracy of an aligner. The mapping between *k*-mer seeds and MEMs (Fig. 1) shows an equivalence of fixed-size seeding and variable-size seeding with respect to such upper bound considerations.

We present and analyze an efficient algorithmic bridge for computing variable-size seeds out of fixed-size seeds. In contrast to previously proposed extend-purge strategies [16-19], we follow a merge-extend strategy that avoids the creation of unnecessary MEM duplicates (Alg. 1, Section 2.4). The detailed differences are discussed in the methods section and the superiority of our approach is shown in the results section. Further, we introduce two seed filters for performantly extracting SMEMs (Alg. 2a, Section. 2.6) and maximal spanning seeds from MEMs (Alg. 2b, Section. 2.6). Additionally, we introduce the notion seed entropy as the number of nucleotides that a single seed contributes to an accurate alignment on average. In the results section, we show that SMEMs have a higher seed entropy than MEMs and that maximal spanning seeds have the highest seed entropy among all three. We investigate the impact of occurrence filtering techniques of state-of-the-art aligners with respect to the mapping and hierarchies shown in Fig. 1. Finally, we present use cases for our algorithmic approaches.

## 2 Materials and Methods

### 2.1 Informal Description

We first introduce our approach informally. Seeds can be visualized by drawing them as 45° lines in a plane that is spanned by reference (x-axis) and query (y-axis). Fig. 2 shows an example for this kind of seed visualization. A detailed description of this representation scheme of seeds can be found in [2]. If *k*-mer seeds are visualized as proposed, some of them overlap or touch one another. Such overlapping or touching seeds can be merged into one single long seed. By performing all possible merges, we get the set of all MEMs. A major disadvantage of *k*-mers is the large size of the required hash-tables. For overcoming this problem, the concept of minimizers was introduced in [20]. Our proposed approach delivers MEMs for minimizers as well if the endpoints of all merged *k*-mers are additionally extended. However, the length of this extension has a strict upper bound. Therefore, the proposed technique delivers long seeds of high entropy out of *k*-mers. The practical value of our technique is an observed runtime superiority regarding index creation and seed computation over the FMD-index as well as a reduction of the number of seeds compared to minimizers.

**Figure 2.**
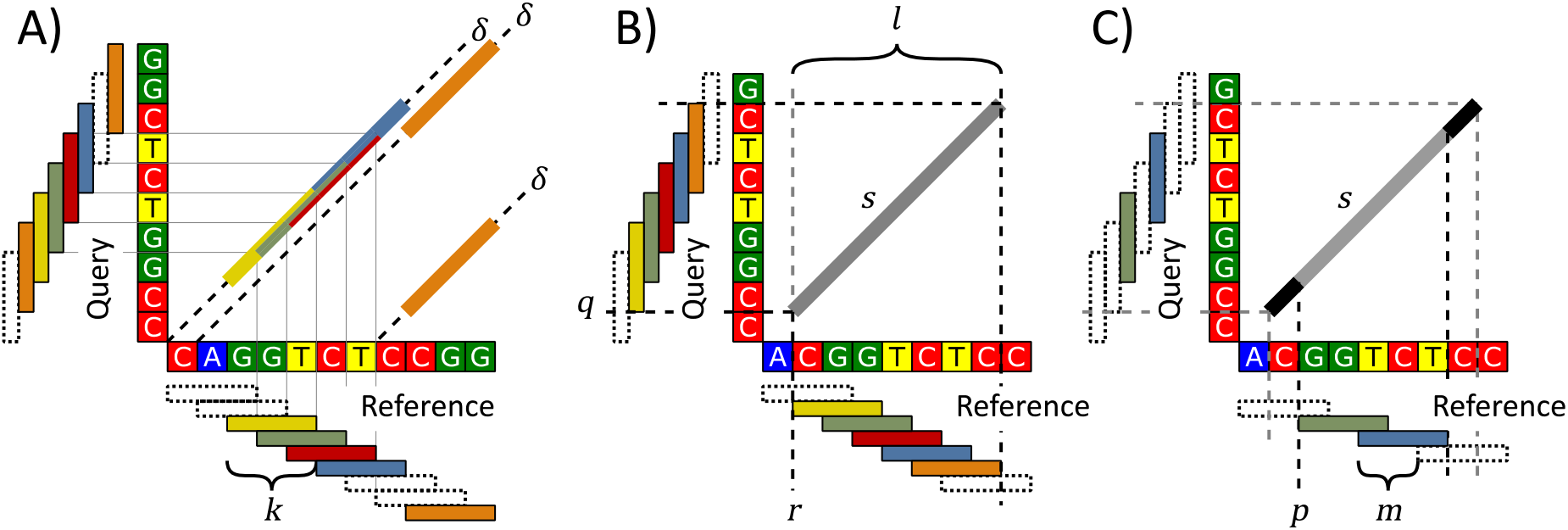
Several aspects of the MEM computation out of *k*-mers. **A)** In a plane spanned by reference (x-axis) and query (y-axis), overlapping *k*-mers form MEMs. The yellow, green, red and blue 3-mers form a single long MEM that spans 6nt. Overlapping *k*-mers are always on the same *δ*-line, which groups seeds according to their *δ*-value. **B, C)** In contrast to plain *k*-mers, *m*-step *k*-mers may merely form subparts of a MEM. The grey MEM *s* = (*q, r, l*) is missing the first and last nucleotide in C), because the yellow and orange 3-mers are not part of the 2-step 3-mers. The black end sections of s require extension for being discovered.

We now present the construction of MEMs out of minimizers. For this purpose, we first describe our technique in the context of *k*-mers with step size of one. Next, we extend the concept to *m*-step *k*-mers, before we finally show that it is applicable for minimizers as well.

### 2.2 Definitions and Notations

Let Σ = {A, C, G, T} be the alphabet and let *W*:= *w*_0_*w*_1_…*w*_*n*−1_ ∈ Σ^+^ be a non-empty word over Σ. A substring *w_x_w*_*x*+1_…*w*_*y*−1_ is denoted by *W*[*x, y*), while a character access to *w_i_* is denoted by *W*[*i*]. {(*x, W*[*x, x* + *k*)): 0 ≤ *x* ≤ |*W*| − *k*} is the set of all positioned *k*-mer sequences over *W* and is denoted by *W^k^*. Let *R, Q* ∈ Σ^+^ be a reference and query sequence, respectively. A seed is a perfectly matching section between *Q* and *R*, which is represented as a triple (*q, r, l*) with 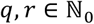, where *q* and *r* are the starting positions on *Q* and *R*, respectively. *l* denotes the length of the seed 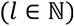. Let *s* = (*q, r, l*) be such a seed. The 5-value of s is defined as *r* − *q*. Using *R*(*s*), we denote the subpart *R*[*r, r* + *l*) of the reference that belongs to *s*. Accordingly, *Q*〈*s*〉 denotes *Q*[*q, q* + *l*). Hence, a seed describes the equivalence of *R*〈*s*〉 and *Q*〈*s*〉.

For a reference *R* and query *Q*, the set of all MEMs is defined, in accordance with [6], as follows: For all pairs of reference-query positions (*x, y*), with *R*[*x*] = *Q*[*y*], 0 ≤ *x* < |*R*|, 0 ≤ *y* < |*Q*|, we get a MEM (*q, r, l*) as follows: Let *i* be the minimal value such that *R*[*x* − *i* − 1] ≠ *Q*[*y* − *i* − 1]. If there is no such value, then *i* = min(*x, y*). Using *i*, we set *q* = *y* − *i* and *r* = *x* − *i*. Further, let *j* be the minimal value such that *R*[*x* + *j*] ≠ *Q*[*y* + *j*]. If there is no such value, then *j* = min(|*R*| − *x*, |*Q*| − *y*). Using *j*, we set *l* = *j* + *i*. The above definition can repeatedly create identical MEMs. We assume that the set is purged of such duplicates.

Given a reference *R*, query *Q*, and their corresponding sets of *k*-mer sequences *R^k^* and *Q^K^*: Let *C^k^*: = {*Q^k^* × *R^k^*} be the cross product of all positioned *k*-mer sequences on *Q* and *R*. Each pair ((*q, W_Q_*), (*r, W_R_*)) ∈ *C^k^* with *W_Q_* = *W_R_* defines a *k*-mer seed (*q, r, k*). The set of all such seeds over *R^k^* and *Q^k^* is denoted by *Seeds*(*Q^k^, R^k^*). We call two seeds overlapping if they have identical δ-values and overlapping or touching reference intervals. Trivially, this implies that overlapping seeds have overlapping query intervals as well. Together, a set of overlapping *k*-mer seeds represents a larger region of equivalence between reference and query (see Fig. 2A). These larger regions correspond to the regions of equivalence described by the set of all MEMs.

### 2.3 Algorithmic Approaches – Getting variable-size seeds from fixed-size seeds

Algorithmically, we can perform seed overlapping as follows: First, for a reference *R* and a query *Q*, we compute *Seeds*(*Q^k^, R^k^*) and store these seeds in an array *K*. Then, we sort *K* ascendingly according to the *δ*-values of all seeds. Seeds of equal delta value are sub-sorted according to their query position. By doing so, we guarantee that the sets of mutually overlapping *k*-mers appear consecutively in *K*. In an iteration over *K*, we merge overlapping *k*-mer seeds. The time complexity of these operations is bounded by the sorting of *K*.

If the proposed algorithmic approach is applied to a set of *k*-mer seeds, it delivers a set of MEMs. Fig. 2A visualizes an example for the computation of three MEMs out of six 3-mer seeds. In the following section, we extend our approach towards minimizers.

### 2.4 Extension of Algorithmic Approach to *m*-step *k*-mers

We first introduce *m*-step *k*-mers. For a word *W* and a given step-size *m*, the set of all positioned *k*-mer sequences (*x*, W[x, x + k)) fulfilling *x* mod *m* = 0 is called *m*-step *k*-mer sequences over *W* and is denoted by 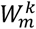. Alg. 1 implements our algorithmic approach in Pseudocode.

We store the *m*-step *k*-mer seeds 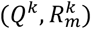 in the array *K*. This altered seed set requires the following two additions to our algorithm: (1) Adjacent *m*-step *k*-mers on a 5-line can be separated by a gap if *m* > *k*. We try to close this gap via an extension process (lines 5 and 6). (2) The ends of a MEM are now not necessarily covered by *k*-mers anymore (see Figure 2C). By iterating diagonally and comparing query and reference for equality, we maximally extend the seeds in *S* (lines 12-16).

#### Algorithm 1. Computation of MEMs out of *k*-mers.

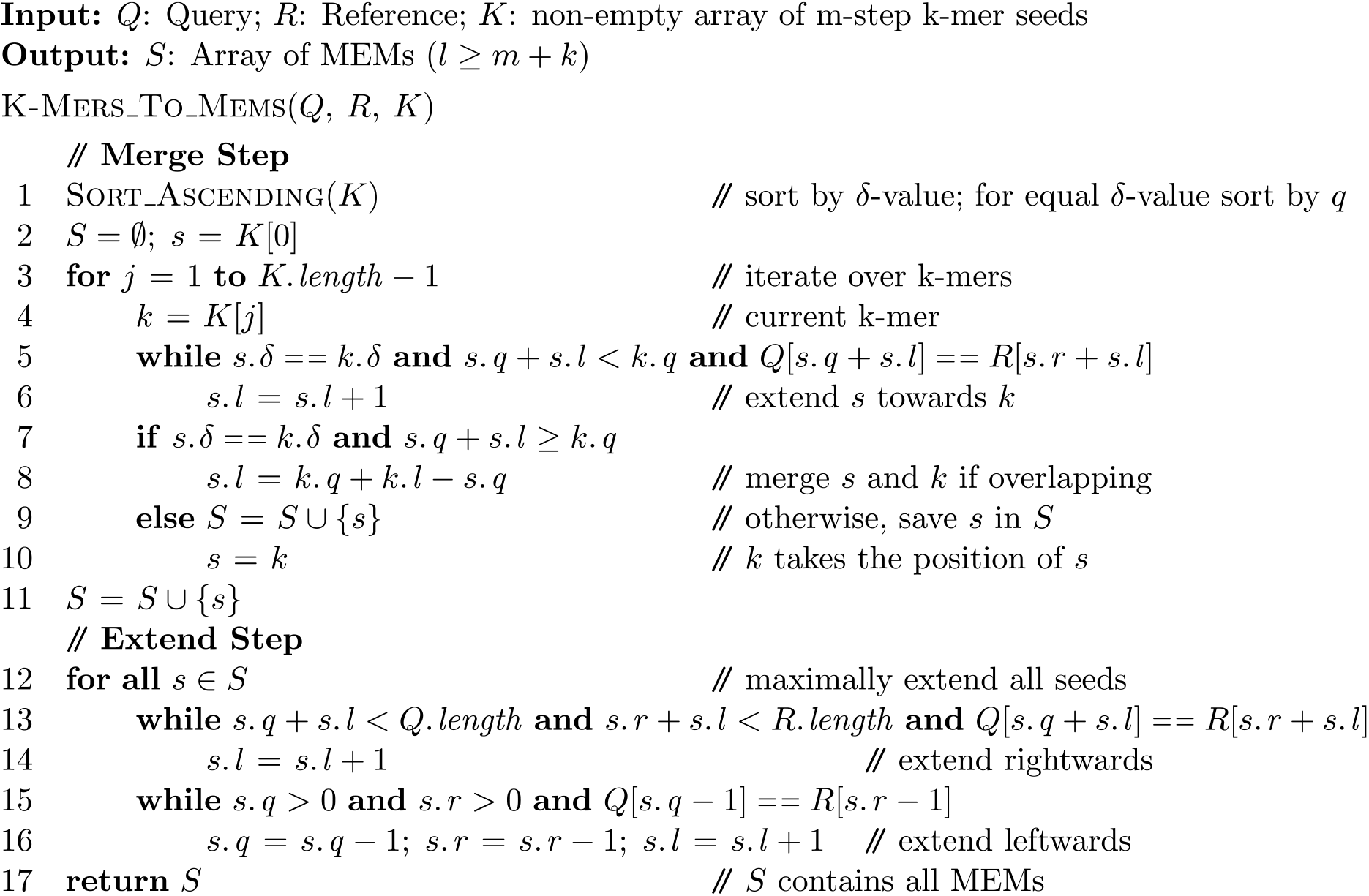

Please note, if we omit the merge step (lines 1-11) and maximally extend all *k*-mers (lines 12-16) immediately, we obtain all MEMs but with the following two significant disadvantages: (1) In the case of a single large match *M* that corresponds to a chain of overlapping *k*-mers, an immediate extension would extend all *k*-mers to the endpoints of *M*. Therefore, the size of *M* determines the amount of extensions required for each *k*-mer of the chain. (E.g. in Fig. 2A, the yellow, green, red and blue 3-mers would all be extended by 3 nt.) (2) A MEM containing multiple *k*-mers would be discovered multiple times. Therefore, we would need an additional purging scheme for getting rid of duplicates.

The above two points matter in the context of the variants of Alg. 1 proposed in [16-19], which all extend *k*-mers before eliminating redundancy. We call their approach extend-purge strategy in contrast to the merge-extend strategy of Alg. 1.

### 2.5 Formal Proof

We now formally prove that our approach successfully retrieves all MEMs out of *m*-step *k*-mers.

#### Theorem 1.

Let *Q* be a query, *R* be a reference and *s* = (*q,r,l*) be a MEM over *Q* and *R*. If *l* ≥ *k* + *m*, then there is at least one *k*-mer seed covered by *s* in *Seeds*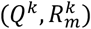.

**Proof.** Let *s*′ = (*q*′, *r*′, *k*) be the sub-seed of *s* with *q*′ = *q* + *x*, *r*′ = *r* + *x* for the offset *x* = *m* − *r* mod *m* and the *k*-mer size *k*. By showing that *s*′ is in *Seeds*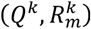, we prove the theorem. Let (*r*′,*R*〈*s*′〉) and (*q*′, *Q*〈*s*′〉) be two positioned *k*-mer sequences. Trivially, (*q*′,*Q*〈*s*′〉) ∈ *Q^k^* and (*r*′,*R*〈*s*′〉) ∈ *R^k^*. We have to show that 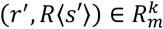, which immediately follows from *r*′ mod *m* = 0. Finally, we have to show *Q*〈*s*′〉 = *R*〈*s*′〉: Due to *s*, the words *R*〈*s*〉 and *Q*〈*s*〉 on reference and query must be equal. Considering *x* ≥ 0 and *x* + *k* ≤ *l*, the subwords *R*〈*s*′〉 = *R*[*r* + *x,r* + *x* + *k*) and *Q*〈*s*′〉 = *Q*[*q* + *x,q* + *x* + *k*) must be equal as well. Altogether, *s*′ results from (*q*′,*Q*〈*s*′〉) and (*r*′,*R*〈*s*′〉).■

#### Theorem 2.

Given a MEM *s* = (*q, r, l*) over *R* and *Q*. For all pairs of seeds *s*′ = (*q*′, *r*′, *k*), *s*′′ = (*q*′′, *r*′, *k*) in 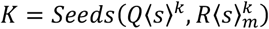, there is a chain of overlapping seeds in *K* connecting *s*′ and *s*′′.

**Proof.** By developing the claimed chain. Without loss of generality, we assume that *r*′ < *r*′′. Let *T* = {(*q* + *m* * *i, r* + *m* * *i, k*) ∈ *K*|*r*′ ≤ *r* + *m* * *i* ≤ *r*′′} be all seeds between *s*′ and *s*′′. The seeds (*q* + *m* * *i, r* + *m* * *i, k*) and (*q* + *m* * (*i* + 1), *r* + *m* * (*i* + 1), *k*) are always overlapping (by *k* − *m* nt) neighbors in *T*. Therefore, *T* represents the claimed chain. ■

#### Theorem 3.

Let *s* = (*q, r, l*) be a MEM over *R* and *Q*. In 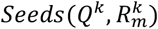, there exists at least one *k*-mer *s*′ = (*q*′, *r*′, *k*) within *m* nt from *r* (i.e. *r* + *m* ≥ *r*′) and at least one *k*-mer *s*′′ = (*q*′′, *r*′′, *k*) within *m* nt of *r* + *l* (i.e. *r*′′ + *k* + *m* ≥ *r* + *l*).

**Proof.** Theorem 1 immediately shows the existence of *s*′. The existence of the *k*-mer *s*′′ can be proven by following the steps of theorem 1 with the modification of choosing *x* as *l* − *k* − (*l* + *r* − *k*) mod *m*. ■

Theorems 2 and 3 prove, that the maximal length that a seed is extended in Alg. 1 is *m* nt. The following corollary follows directly from Theorem 1 and 2:

**Corollary 1.** For a reference *R* and a query *Q* and the *k*-mer seed set 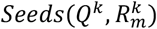. Alg 1. computes all MEMs with a size ≥ *m* + *k* exactly once.

### 2.6 Extension to Minimizers and Extraction of SMEMs and Maximal Spanning Seeds

The proposed algorithmic approach for *m*-step *k*-mers can be applied to minimizers with a window of *m*, because the distance between two adjacent (*m, k*)-minimizers is always ≤ *m* (as proven in [20]).

#### Algorithm 2a. Computation of SMEMs out of MEMs.

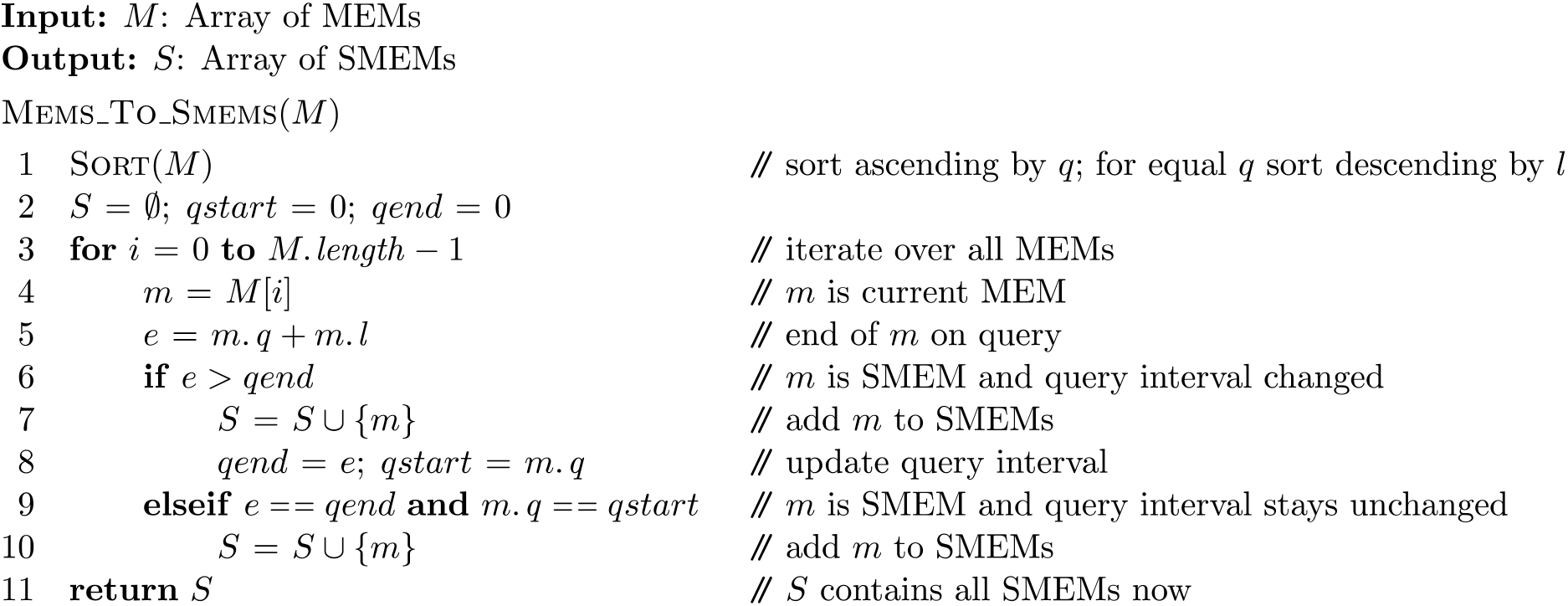

Algorithm 2a shows the Pseudocode for the extraction of SMEMs out of MEMs using a sorting followed by a single sweep over all MEMs. In order to prove the correctness of the algorithm, we characterize SMEMs (in accordance to [6, 15]) as follows: A SMEM is a MEM that is not enclosed by another MEM on the query. The above algorithm identifies all enclosed MEMs by iterating over them ordered by their start positions on the query.

The variable *qend* keeps the rightmost end-position of all MEMs visited during the iteration so far. For each iteration of the loop, we have: If the current seed *m* ends before *qend*, then *m* cannot be a SMEM, because the previously visited MEM that assigned *qend* encloses *m. qstart* memorizes the start position of the last SMEM. If start and end of the previous SMEM are equal to the current SMEMs start and end (line 9), then the current seed is a SMEM (line 10) that spans the same query interval as the previous one but has a different reference interval. Alg. 2a requires time *O*(*n* log *n*) for a set of *n* MEMs, assuming the initial sorting is done in time *O*(*n* log *n*).

#### Algorithm 2b. Computation of Maximal Spanning Seeds out of MEMs.

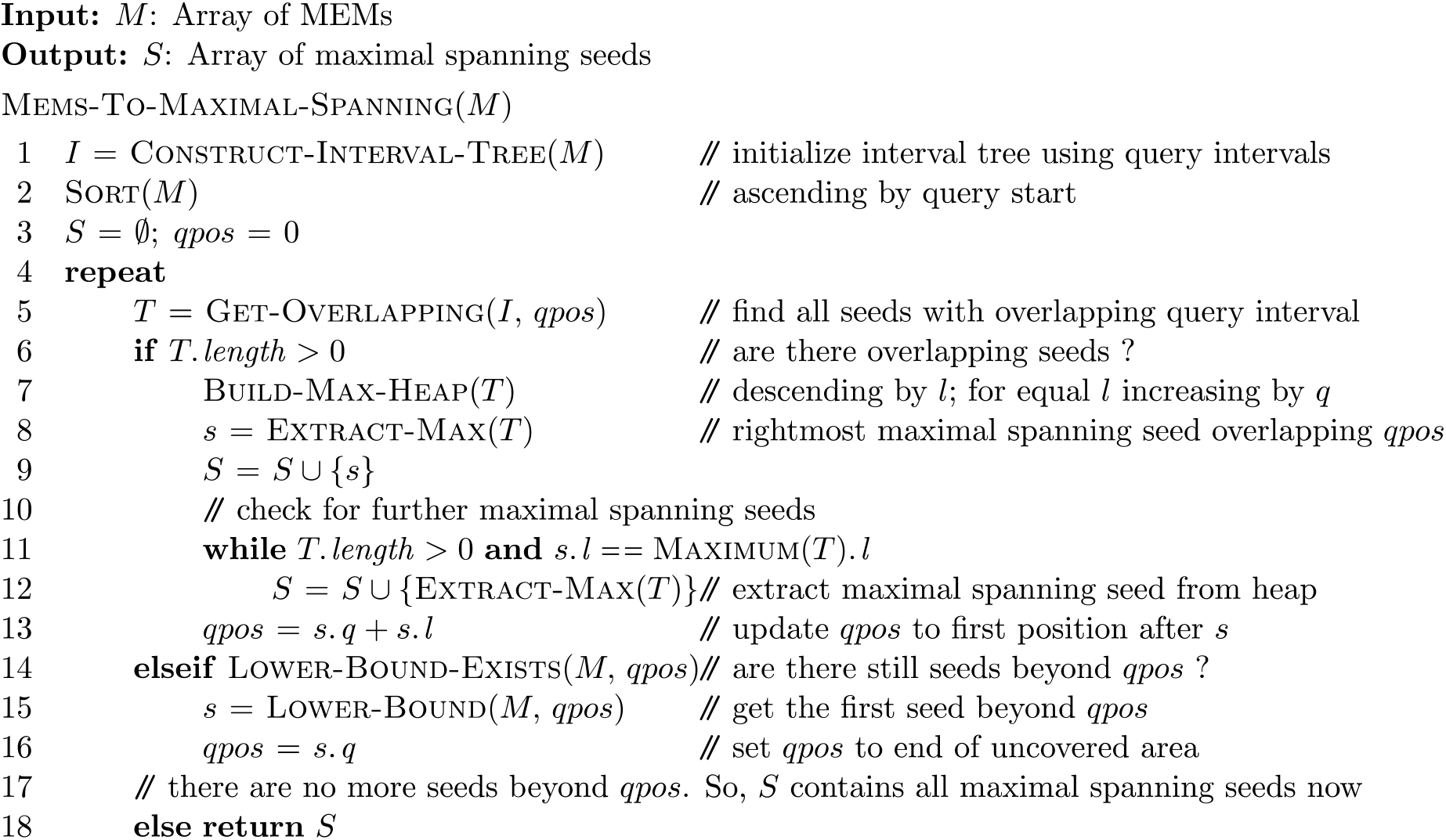

Within the MEMs, we identify the set of maximal spanning seeds as follows: A MEM is a maximal spanning seed if and only if it comprises at least one query position, where it is not covered by another longer MEM. Please note that the maximal spanning seeds always represent a subset of the SMEMs. (A detailed comparison of SMEMs and maximal spanning seeds can be found in [2].)

We now informally describe the approach of Alg. 2b. A detailed analysis is given in Supplementary Note 1. In the central loop (lines 4-18), we visit all maximal spanning seeds exactly once via *qpos*. In each iteration, we distinguish between two situations:

1. *qpos* is within a set of seeds (Boolean condition in line 6 is true). We identify all maximal spanning seeds (line 8-12) by building a heap according to the seeds’ lengths (line 7) for all seeds overlapping *qpos* (line 5). If there are several maximal spanning seeds at *qpos*, then they are all identified in the while-loop (lines 11 and 12). Line 13 updates *qpos* for the next iteration. This guarantees that no maximal spanning seed is discovered twice or lost.
2. *qpos* is at the beginning of an area without seeds at all (Boolean condition in line 6 is false). In this case we move *qpos* to the first seed after this area (lines 14-16) or we return the computed set *S*, because there are no unprocessed seeds anymore (line 18). If *qpos* is moved to the start of a seed, we continue with case 1 in the next iteration.

#### Theorem 4.

For a set of *n* MEMs, Alg. 2b) requires time *O*(*n* log *n*), assuming the sorting (line 2) and interval tree construction (line 1) happen in time *O*(*n* log *n*).

**Proof.** The central loop (lines 4-18) is executed ≤ 2*n* times. Further, lines 5, 14 and 15 require time O(logn) per iteration. In line 5, the accumulative size of all *T* is ≤ *n*, because each MEM can only be part of a single iteration’s *T*. Therefore, all operations performed on *T* (lines 7, 8, 11 and 12) require an accumulative time of *O*(*n* log *n*). ■

## 3 Results

We now prove the practical value of the proposed merge-extent technique by comparing its runtime behavior with the extend-purge strategy as well as FMD-index based seeding. Further, we introduce and discuss seed entropy. Finally, we report about the effects of occurrence filtering in the context of seeding and present some practical use cases of our approach.

Our benchmarking relies on simulated PacBio reads, where the error profiles are sampled from the HG002 GIAB dataset [21]. The simulation scheme is described in Supplementary Note 2. In the diagrams, we compare the behavior of seeding techniques for various error rates, where we start with perfect reads and increase the error rate until we meet the error profile of a specific kind of sequences reads. An algorithmic description of the used extend-purge strategy that is in accordance with the approaches presented in [16-19], is given in Supplementary Note 3.

Because the superiority of the merge-extend strategy increases with increasing read length, our analysis focuses on long reads. However, Supplementary Note 4 contains an analysis for Illumina reads as well.

### 3.1 Time Evaluation

We now analyze the runtime behavior of seeding techniques as visualized in Fig. 3. The claimed theoretical superiority of our merge-extend strategy is reflected by the runtimes. The superiority becomes particularly well visible for high quality reads. Due to the expected improvements regarding the quality of long reads, these differences are of increasing relevance in practice. Further, the benchmarking shows that the FMD-index is not well suited for the computation of MEMs. Here, the runtimes grow almost exponentially with increasing read quality. For computing MEMs, we adopted the algorithm presented by Ohlebusch et al. [22].

**Figure 3.**
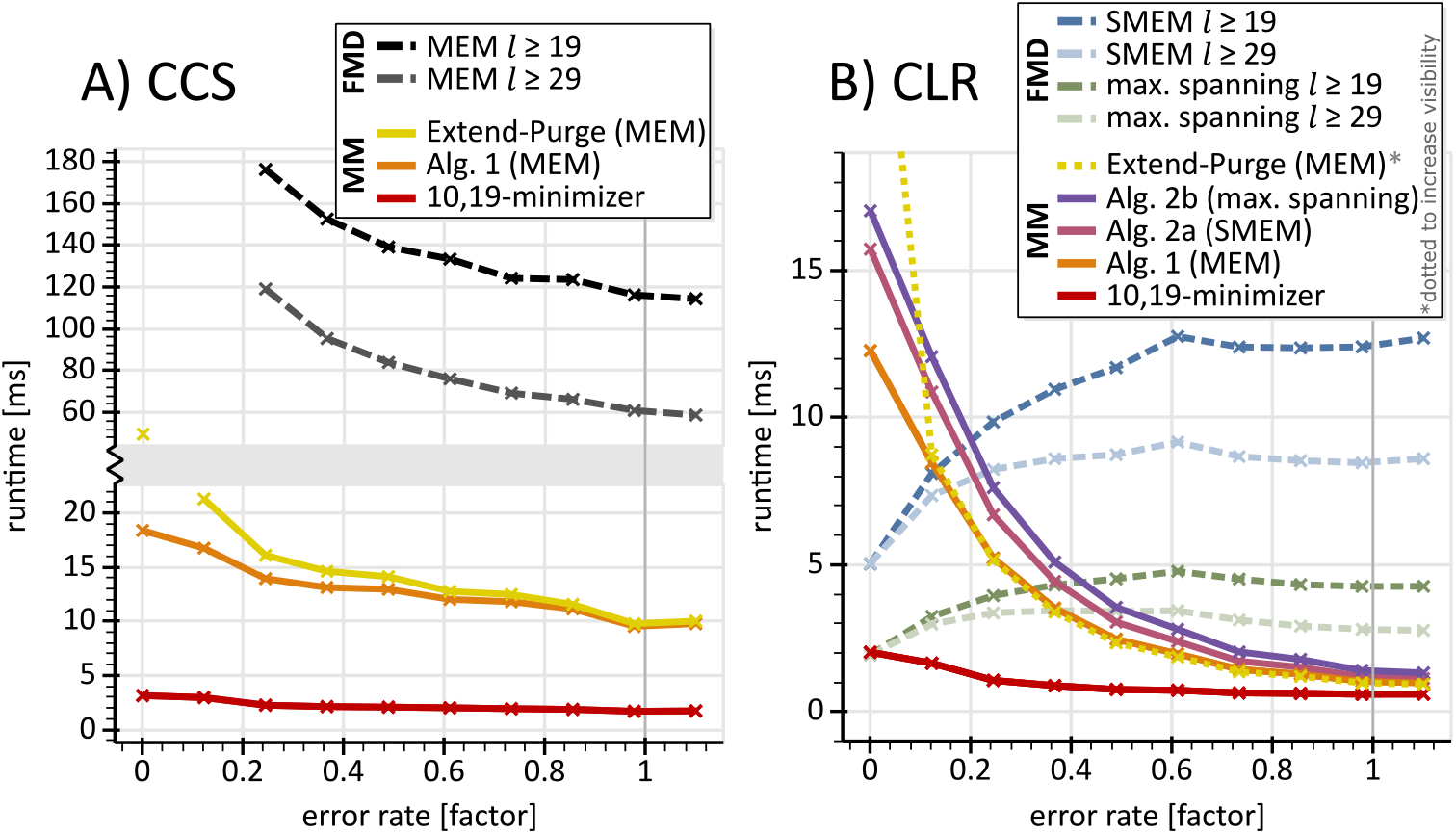
Time evaluation. **A)** shows the runtimes for circular consensus sequencing (CCS) PacBio reads with a focus on MEM computation: The x-axis denotes a factor applied to the error profile of CCS PacBio reads, where zero corresponds to error free reads and one corresponds to reads following the measured error profile. The detailed scheme used for error rate computation is described in Supplementary Note 2. The generation of MEMs via the FMD-index is unreasonably slow compared with minimizer based approaches. Our merge-extend strategy displays a general superiority over the extend-purge approach, where the superiority grows with increasing read quality. For reads of high quality, the extend-purge strategy tends to extend multiple minimizers to the same MEM before purging all duplicates. In contrast, our merge-extend strategy computes each MEM only once. This problem also worsens with increasing read length because the extensions increase in size. **B)** shows runtimes for continuous long PacBio reads (CLR). Here, we additionally show runtimes for the SMEM and maximal spanning seed computation. The opposing behavior of the curves for the FMD-index (dashed lines) and corresponding curves for minimizers (solid lines) indicates a superiority of the latter approach for state of the art reads. The pink and purple graphs include the time required for previous steps, i.e. Alg. 1 and minimizer computation. Accordingly, the orange graph includes the time required for minimizer computation.

Fig. 4B proves the practical value of our approach for filtering SMEMs and maximal spanning seeds out of MEMs (Alg. 2A and 2B). Here, the runtimes of seeding correlate with the size of the computed seed sets. The more seeds in a set, the longer it takes to compute it. With increasing read quality, the number of SMEMs and maximal spanning seeds decreases while the number of MEMs and minimizers increases. Therefore, the direct computation of SMEMs and maximal spanning seeds becomes faster with increasing read quality, while the extraction of these seeds from a set of MEMs becomes slower. The FMD-index allows such a direct computation of seeds, while Alg. 2a and Alg. 2b extract them from the set of MEMs. This explains the reciprocal behavior of the dashed and solid curves in Fig. 3B. Thus, for SMEM and maximal spanning seed computation, the preferred algorithmic approach depends on the quality of the reads. For continuous long PacBio reads (CLR), our measurements indicate that Alg. 2a and Alg. 2b outperform the FMD-index.

**Figure 4.**
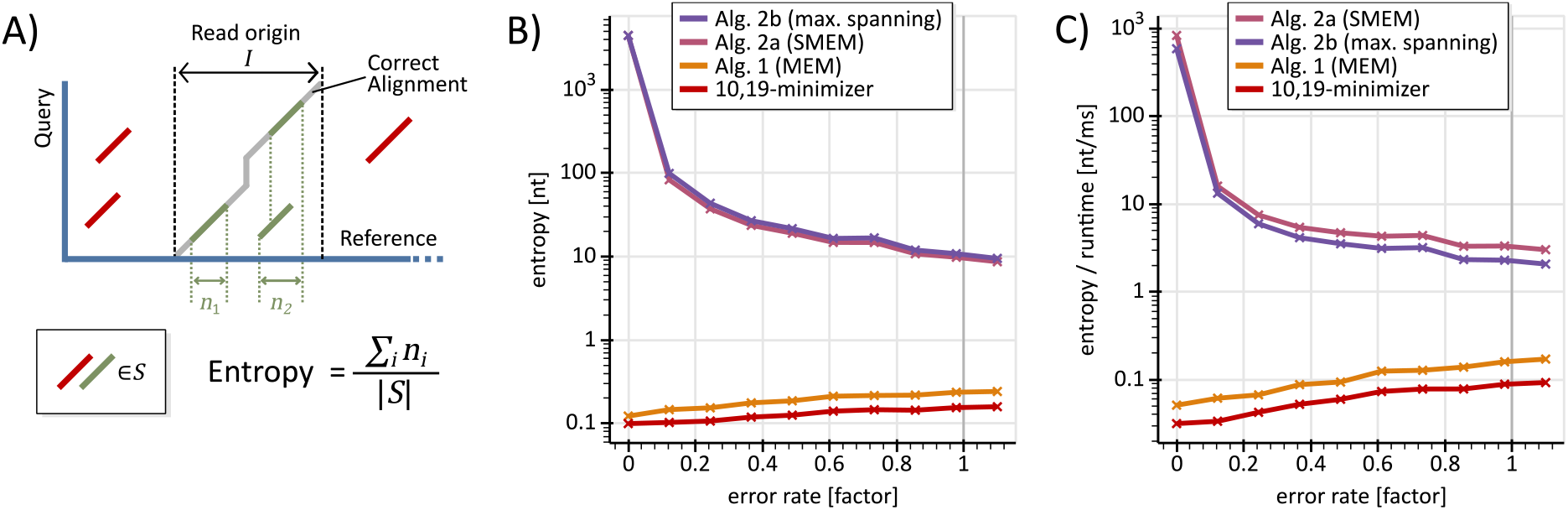
Seed entropy analysis. **A)** Informally, for a read *r*, the entropy measures the ratio between the size of the reference area covered by “correctly placed” seeds of *r* and the number of seeds for *r*. **B)** analyzes the seed entropy for CCS PacBio reads according to the entropy computation given in A). Each point in the diagram shows an average value for 1,000 reads. The red curve displays the behavior of unprocessed minimizers, while all other curves show benchmarks for the proposed algorithmic approaches. **C)** visualizes the ratio entropy over runtime, where the benchmarking environment is equal to B).

Although the extraction of maximal spanning seeds (purple curve) requires an interval tree, a heap and an initial sorting operation, it is only slightly slower than the SMEM filtering (pink curve). This can be explained by different behaviors of Alg. 2a and 2b in their central loops, where Alg. 2b merely iterates over maximal spanning seeds (and gaps among them) while Alg. 2a iterates over all MEMs.

### 3.2 Seed Entropy Analysis

We now define the notion seed entropy and analyze its behavior:

#### Definition

Let *Q* be a read that originates from a reference interval *I*. Let *S* be a set of seeds for *Q*. The seed entropy of *S* for *Q* is the ratio *n*/|*S*|, where *n* is the number of nucleotides in *I* that are covered by seeds of *S* and |*S*| is the size of the set *S*.

The entropy expresses the number of nucleotides that a single seed contributes to an accurate alignment on average. Expressed in other terms, the entropy is the ratio of correct data over incorrect data with respect to seeds.

Within the interval *I*, the seed entropy does not distinguish between seeds that contribute to an accurate alignment and seeds that do not. We call contributing seeds correct and non-contributing seeds incorrect. Incorrect seeds can erroneously increase the coverage *n*. A more accurate definition of seed entropy could be established on the foundation of correct seeds merely. However, Survivor [23], the read generator used here, does not deliver sufficient information for identifying correct seeds in *I* and so we have to rely on the proposed definition. Fig. 4 visualizes our entropy measurements for several seed sets using the various algorithmic approaches proposed here. The environment used for benchmarking is described in Supplementary Note 2.

The curve for minimizers is always below the curve for MEMs (computed by Alg. 1). This can be explained as follows: Combining seeds (Alg. 1, lines 1-11) reduces the number of seeds while maintaining the coverage. Additionally, extending on both ends (Alg. 1, lines 12-16) increases the coverage while keeping the number of seeds. Hence, Alg. 1 can only increase the seed entropy. Diagram 4B shows that the entropy for SMEMs is always significantly higher than for MEMs. This confirms that SMEMs are a cleverly chosen subset of MEMs that is well suitable for alignments. Maximal spanning seeds always have the highest entropy among all four seed sets. The ratio entropy over runtime (Fig. 4C) indicates that the tradeoff between additional runtime and increased entropy is in favor of Alg. 2a and Alg. 2b over MEMs and minimizers.

Our measurement express the entropy of a seed set and not its capability to allow accurate alignments. The proposed algorithmic approaches can only deliver seeds that are discoverable via minimizers and nothing beyond.

### 3.3 Filter and Effects

Most aligners (e.g. Minimap 2 [3], MA [2], BWA-MEM [4]) use a filtering scheme in order to cope with repetitive regions of the genome. Using a threshold, seeds of query intervals that show an excessive amount of occurrences (i.e. a high ambiguity) on the reference are purged during seeding. However, minimizers and FMD-index apply these filters on different stages of the seeding process. The FMD-Index filters after the completion of the extension (using the size of the suffix-array intervals), while minimizers filter using the size of the hash-table buckets. In Fig. 5, we evaluate the effect of this difference in the context of the human genome.

**Figure 5.**
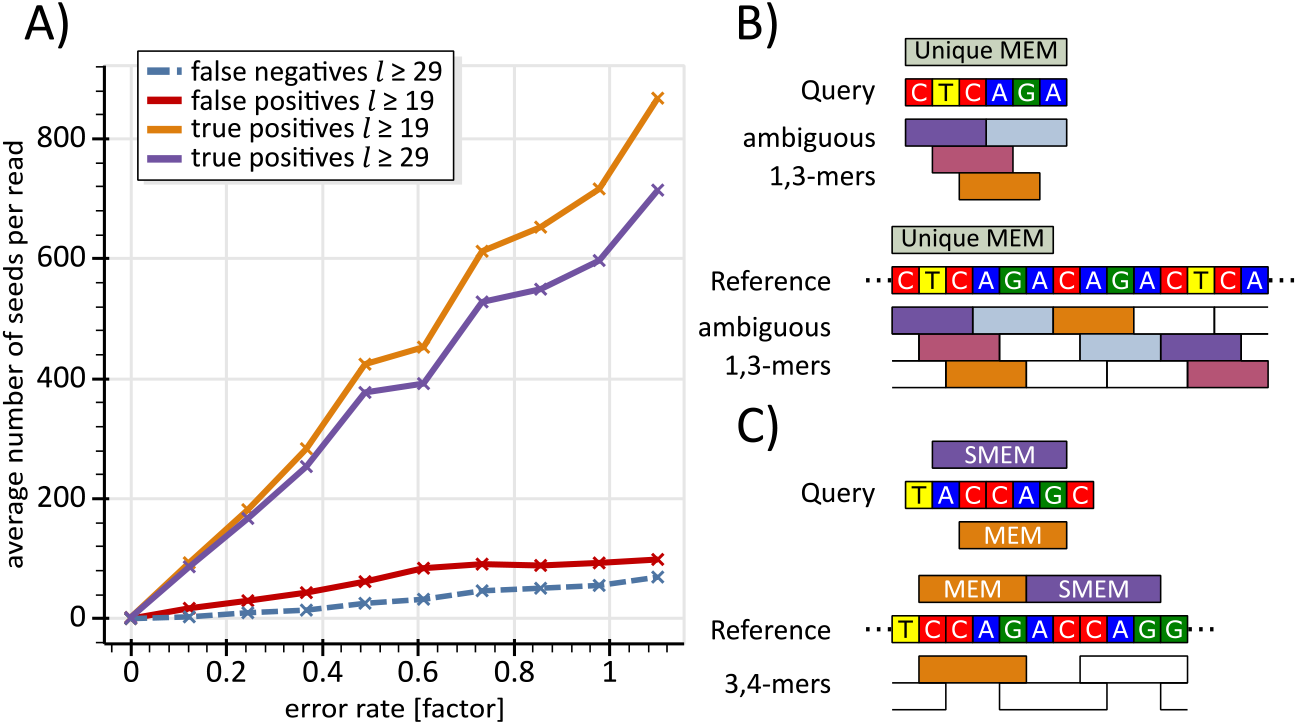
Analysis and effects of occurrence filtering. **A)** The four curves visualize the behavior of occurrence filters in the context of the computation of SMEMs via Alg. 2A and the FMD-index. The x-axis represents the error rate of reads as described in Fig 3. Each dot shows the average value for 1000 CCS PacBio reads generated using Survivor [23]. The red curve shows the average amount of seeds generated by Alg. 2a that do not exist in the set of SMEMs (with *l* ≥ 19) computed by the FMD-index. The blue curve displays the opposite information; it shows all seeds (with *l* ≥ 29) computed by the FMD-Index that do not appear in the seed set computed by Alg. 2a. The orange and purple curves inform about the number of seeds shared by Alg. 2a and the FMD-index with *l* ≥ 19 and *l* ≥ 29, respectively. **B)** There is a MEM covering the unique sequence CTCAGA on query and reference. Assuming, the occurrence threshold for minimizers is set to two, the four colored 1,3-mers are purged. In this case, the MEM cannot be discovered by Alg. 2a. Using the FMD-index, this MEM is discovered directlyand not purged by the occurrence filter, since it occurs only once. **C)** explains the generation of SMEMs that are false positives. The purple SMEM cannot be discovered using the 3,4-mers, since no 3,4-mer is contained within the seed. However, the orange MEM can be discovered using the orange 3,4-mer. Now, Alg. 2a will not delete this MEM, since it is not enclosed by any other seed.

In order to compare FMD-index based SMEMs with minimizer based SMEMs in practice, we rely on the following filter settings: For computing minimizers, we use Minimap 2, where ‘-*f*’ is set to zero and the *‘--min-occ-floor’* threshold is set to 200. For the FMD-Index, MA’s *‘Maximal Ambiguity’* parameter is set to 200. Therefore, both seeding schemes drop seeds occurring more than 200 times on the reference.

Without filtering, the theoretical equivalence implies that the red and blue curves of Fig. 5A must constantly be zero. The blue line, SMEMs found via the FMD-index merely, results from situations like the one depicted in Fig. 5B: There exist 19-mers that exceed the occurrence threshold while the respective maximally extended seed does not. However, as shown in Supplementary Note 5, such seeds have a low entropy and so they are not expected to contribute to an accurate alignment.

Further, there are SMEMs that are found via Alg. 2a but not via the FMD-index. These SMEMs are false positives and their appearance is explained in Fig. 5C: Due to the absence of minimizers, a SMEM stays undiscovered. Instead, the largest MEM inside the query interval of that SMEM appears as a false positive SMEM now. These false positives have a very low entropy as shown in Supplementary Note 5. Please note, without filtering and for *l* ≥ *w* + *k* (for *w, k*-minimzers) such false positives vanish since the absence of minimizers becomes impossible.

Supplementary Note 5 contains a corresponding occurrence filter analysis for maximal spanning seeds as well as CLR PacBio reads.

### 3.4 Single-use Indices as Application of the Proposed Approach

The hash tables used for minimizer seeding can be computed in short time by a single scan over the reference. Therefore, minimizer indices are well suited as “single-use” indices for subsections of the human genome. We now describe two application scenarios for single-use indices:

Scenario 1: For filling gaps between seeds, aligners often rely on Dynamic Programming [1-3, 12]. Technically, it is common to start from one seed’s endpoint and to find a path to the next seed’s start point throughout the matrix. There are structural variants that cannot be detected via such a path-oriented strategy. Examples of such structural variants are micro duplications and micro translocations as e.g. shown in Fig. 6. Dynamic programming runs into trouble there. It cannot locate the green and orange matches, because they are positioned above and below the band, respectively. Increasing the bandwidth is of limited use, since backtracking can only incorporate one of both seeds into an alignment’s path. Further, an index over the whole genome would also miss these seeds due to their size. However, they can be discovered by computing a single-use index that spans the Dynamic Programming area.

**Figure 6.**
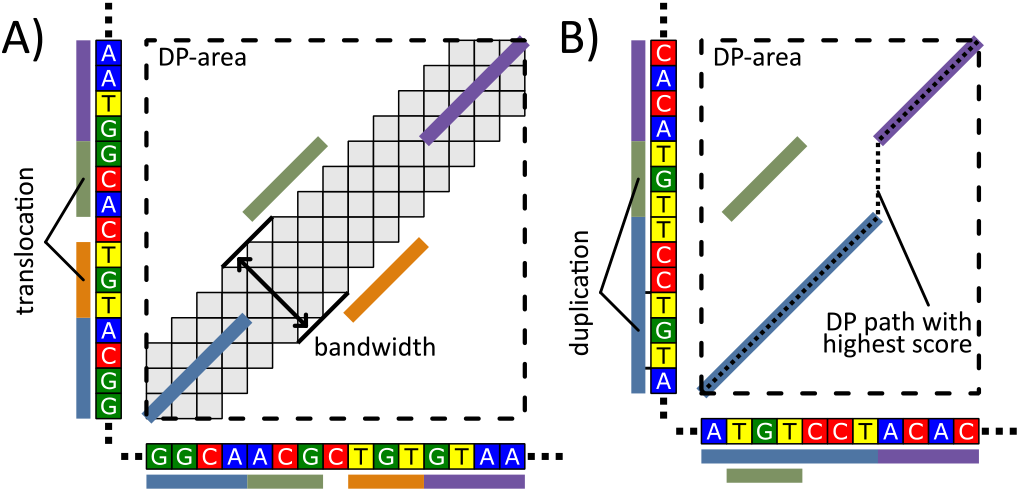
Dynamic programming (DP) encounters issues recognizing the two structural variants shown above. However, both variants could be discovered by reseeding in the DP area. **A)** shows a micro-translocation. DP can discover neither the green nor the orange seed due to their out-of-band locations. Unbanded DP could discover one of both seeds merely, but never both. **B)** shows a micro-duplication. Neither banded nor unbanded DP can discover the green seed. Instead, a pseudo insertion would be reported between the endpoints of the blue and purple seed.

Scenario 2: Due to the repetitiveness of genome sections, the initial seeding of a read might deliver a candidate region merely without precisely placing the read within such a region (e.g. a read of a tandem repeat). In this case, a reseeding with smaller seeds in the candidate region can help acquiring a more accurate alignment. As in scenario 1, indices for minimizers allow such a reseeding in short time and enable the fast delivery of MEMs, SMEMs and maximal spanning seeds via Alg. 1, Alg. 2a and 2b, respectively.

## 4 Discussion

Alg. 1 represents a surjective and not injective mapping from *k*-mers to MEMs (see Fig. 1). Therefore, informally spoken, everything that can be discovered with *k*-mers (fixed-size seeding) can also be discovered using MEMs (variable-size seeding) and vice versa. This implies that there is no theoretical superiority of one of these seeding techniques in contrast to assumptions made by us [2] and others [3]. A sophisticated chaining of minimizers should deliver the same alignment accuracy as a sophisticated chaining of MEMs.

Let |*S*|_⊤_ and |*S*|_⊥_ be the absolute number of correct and incorrect seeds in a set of seeds *S*, respectively. Then, we get the following hierarchies:

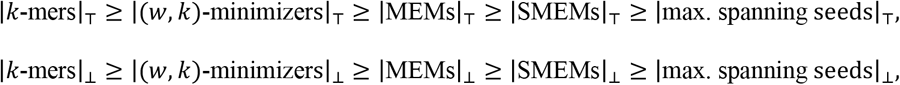

where MEMs, SMEMs and maximal spanning seeds are of size ≥ *w* + *k*. However, our benchmarking suggests a reverse hierarchy regarding the seed entropy (see Section 3.2). Hence, with maximal spanning seeds, an aligner will use the seed set with the highest entropy, but it is at high risk of missing an accurate alignment due to the lack of correct seeds. In contrast, with *k*-mers an aligner has access to the largest number of correct seeds but struggles to cope with the excessive amount of incorrect seeds.

Naturally, each approach that processes a set of seeds (e.g. chaining, clustering) has a time complexity driven by the number of seeds to be processed (e.g. non-heuristic chaining can have a squared worst case complexity [3]). Therefore, the above hierarchies for the number of correct (and incorrect) seeds within sets of seeds also imply hierarchies for the times required to process these sets (processing *k*-mers is the most expensive, while processing maximally spanning seeds is the cheapest). If the runtime of one of our algorithms is less than the time required to process the respectively larger seed set (e.g. from minimizers to MEMs with Alg. 1), we get an overall runtime advantage. Here the accuracy can suffer with SMEMs (Alg. 2a) and maximal spanning seeds (Alg. 2b), since the amount of correct seeds is reduced, as mentioned above.

If a seed processing algorithm can deliver all seeds sorted by their 5-values, the sorting operation of Alg. 1 (line 1, sorting by ó-value) is not required anymore. This gives Alg. 1 a linear runtime complexity and so all MEMs can be computed in linear time. An example of a seed processing strategy that can exploit this aspect of Alg. 1 is the Strip of Consideration proposed in [2].

FMD-Index computation can be boosted by exploiting concurrency. Trivially, such introduction of concurrency is possible in the context of minimizers as well. We limit our benchmarking to a single thread in order to get a fair comparison of both techniques.

Our analysis bases on the human genome and a limited range of parameter combinations. A more comprehensive study could yield more insights here. Further, the design of more sophisticated heuristics for occurrence filtering of minimizers deserves additional research. Finally, the single-use index approach mentioned in section 3.4 lacks practical evaluation so far.

## 5 Conclusion

Our novel algorithmic approaches allow the generation of SMEMs and maximal spanning seeds using minimizers. Particularly for long reads of high quality, our merge-extend strategy for MEM computation is faster than existing extend-purge approaches. In the context of aligner design, the proposed hierarchies within fixed-size seeding and variable-size seeding together with their respective entropies can be used for choosing an appropriate seeding technique (e.g. choosing SMEMs over MEMs is expected to decrease runtime but for the price of a slightly worse accuracy). The reported impact of occurrence filters helps assessing their effects with respect to the accuracy runtime tradeoff of alignments. Summarily, our presented algorithms and insights are valuable in the context of designing and using aligners.

## Abbreviations

CCS: Circular consensus sequencing
CLR: Continuous long read
DP: Dynamic programming
MEM: Maximal exact match
SMEM: Supermaximal exact match

## Declarations

### Ethics approval and consent to participate

Not applicable.

### Consent for publication

Not applicable.

### Availability of data and materials

All code and datasets supporting the conclusions of this article are available via the GitHub repository, https://github.com/ITBE-Lab/seed-evaluation under the MIT License. The evaluation tools are realized using Python 3.6, where time critical components (minimizer computation, proposed algorithms etc.) are implemented in C++ 17. All code runs under Debian Linux 9.12 (stretch).

### Authors’ contributions

The project was conceived by all authors. A.K. and M.S. devised and analyzed all algorithmic schemes with P.S.K. contributing valuable remarks. A.K. and M.S. implemented and conducted all experiments. The manuscript was written by and A.K. and M.S. with assistance of P.S.K.

