## Supplementary Notes for "A performant bridge between fixed-size and variable-size seeding"

### Supplementary Information

#### Supplementary Note 1 - Detailed analysis of Alg. 2b.

**Lemma 5:** Let $M$ and $S$ be the set of all MEMs and all maximal spanning seeds over a given reference and query, respectively. Alg. 2b computes $S$ out of $M$.

**Proof:**

We show the correctness of the following loop invariant for all iterations of the central loop (lines 4 -18):

**Loop invariant:** At position $qpos$, we collect all maximal spanning seeds that encompass $qpos$. $S$ contains all max. spanning seeds that end before $qpos$. There is no seed that occurs twice in $S$.

**Initialization:** For $qpos = 0$, we have $S = \emptyset$. ($S$ is empty, because it comprises all seeds left of $qpos$ and there cannot be a seed ending before position 0.)

**Maintenance:** We have to distinguish 2 cases at $qpos$:

1. No seed overlaps $qpos$: In this case, there is no seed to be collected; $S$ stays unchanged. Lines 14-16 move $qpos$ to the start of the next seed. (The next following case must be case 2.)
2. There are seeds overlapping $qpos$: Let $O$ be the set of those seeds. $O$ is computed in line 5. By definition, a seed is maximal spanning if and only if it comprises at least one query position, where it is not covered by another longer MEM. At position $qpos$, this condition holds for all seeds of $O$ that are added to $S$ in lines 7-12 via the priority queue (max heap). Hence, we collect all max. spanning seeds overlapping $qpos$.
   We now have to show that the next stop of $qpos$ is chosen so that no max. spanning seed is selected more than once (a) or skipped (b):
   Let $s \coloneqq (q, r, l)$ be the seed extracted in line 8.
   a. Due to the heap ordering, $s$ is the rightmost extending seed among all max. spanning seeds in $O$. By setting $qpos$ to the first position after $s$ (line 13), we will never collect any seed twice.
   b. By contradiction, we prove that no max. spanning seed is skipped: Assume these is a max. spanning seed $s' \coloneqq (q', r', l')$ that is skipped, i.e. $s'$overlaps neither $qpos$ nor $q + l$ (the position of $qpos$ in the next iteration). Hence, $qpos < q'$ and $q' + l' \leq q + l$. Since we extracted $s$ at the position $qpos$, we have $q \leq qpos$. Therefore $s'$ is fully enclosed by $s$ ($q < q' < q' + l' \leq q + l$) and cannot be a max. spanning seed.

**Termination:** $qpos$ is increased in every iteration by at least one nt. As soon as $qpos$ is past the end of the query, there cannot be any overlapping seed or any seed to the right of $qpos$. Then, line 18 terminates the central loop.

#### Supplementary Note 2 - Detailed description of the read simulation and error rate

All benchmarking is done using the human reference genome GRCh38.p12 (GenBank assembly accession: GCA_000001405.27). For the simulation of PacBio circular consensus sequence (CCS) reads and continuous long sequence (CLR) reads, we use the program Survivor [1], version v1.0.5-14-g18bf070. For the simulation of 250nt Illumina reads, we rely on the program DWGSIM [2], version 0.1.12-2-g39a1bbb. All diagrams denoting an "error rate" on the x-axis are computed as follows:

At the x-axis position labeled '1', we show measurements for the "standard error rate" of the benchmarked type of reads (PacBio CCS etc.). The standard error rate is chosen as follows:

- For Survivor CLR PacBio reads, we use the error profile provided in the GitHub repository of Survivor.
- For Survivor CCS PacBio reads, the profile is measured using Survivor and the CCS 10kb PacBio reads of the HG002 individual in the GIAB dataset. (ftp://ftp-trace.ncbi.nlm.nih.gov/giab/ftp/data/AshkenazimTrio/HG002_NA24385_son/PacBio_CCS_10kb/)
- All Illumina reads are created using the standard setting of DWGSIM.

At the x-axis position labeled '0', error free reads are used for benchmarking. The fractional values on the x-axis denote factors that are applied to the standard error rate. For values smaller than one, the error rate is decreased; otherwise, the error rate is increased. In more detail, this is done as follows:

- DWGSIM: The modulation of the error rate happens by multiplying the x-axis factor with 0.001 (default value of '-r' rate of mutations parameter) and using the result as the new rate of mutations.
- Survivor: Using the error profile for the standard error rate, we create a specific error profile for a given factor $f$ as follows: In Survivor, error profiles are represented using tables consisting of 6 columns:
    1. a position value $p$
    2. probability $P_{stop}(p)$ that a read ends at $p$
    3. probability $P_{match}(p)$ for a match at $p$
    4. probability $P_{mismatch}(p)$ for a mismatch at $p$
    5. probability $P_{ins}(p)$ for an insertion at $p$

6. probability $P_{del}(p)$ for a deletion at $p$

For each row, we compute factor specific values $P'(p)$ as follows:

$P'_{stop}(p) = P_{stop}(p)$

$P'_{match}(p) = 1 - (1 - P_{match}(p) * f)$

$P'_{mismatch}(p) = P_{mismatch}(p) * f$

$P'_{ins}(p) = P_{ins}(p) * f$

$P'_{del}(p) = P_{del}(p) * f$

All benchmarking with respect to the FMD-index is done using the FMD-index implementation of MA (version 1.1.1-a7a0989) [3]. Computing MEMs using the FMD-index is done via an adoption of the algorithm in [4] from the FM-index to the FMD-index. The generation of minimizers is measured using code of Minimap 2 (version 2.12-r829) [5].

In the context of our benchmarking for seeding, we measure the time required for the actual seed production. For minimizers, this includes the time required for the minimizer computation as well as the time required for the hash table lookup of all reference positions. For the FMD-index, this includes the time required for the extension as well as the time required to extract the reference positions of seeds form the suffix array.

All benchmarking is done on an *AMD Ryzen Threadripper 1950X 16-Core Processor* with 128 GB RAM. For compilation, we rely on gcc (version 6.3.0). As software environment, we use Debian GNU/Linux with a 4.9.0 kernel.

All code is available as open source at: https://github.com/ITBE-Lab/seed-evaluation

#### Supplementary Note 3 – Detailed description of the extend-purge scheme

**Input:** $Q$: Query; $R$: Reference; $K$: non-empty array of m-step k-mer seeds
**Output:** $S$: Array of MEMs ($l \geq m + k$)

```
EXTEND_PURGE(Q, R, K)
    // Extend Step
 1  for all s ∈ K                                   // maximally extend all seeds
 2      while s.q + s.l < Q.length and s.r + s.l < R.length and Q[s.q + s.l] == R[s.r + s.l]
 3          s.l = s.l + 1                           // extend rightwards
 4      while s.q > 0 and s.r > 0 and Q[s.q − 1] == R[s.r − 1]
 5          s.q = s.q − 1; s.r = s.r − 1; s.l = s.l + 1   // extend leftwards
    // Purge Step
 6  SORT(S)                                         // sort by any absolute order among seeds
 7  S = ∅; s = K[0]
 8  for j = 1 to K.length − 1                       // iterate over MEMs
 9      k = K[j]                                    // current MEM
10      if s.q ≠ k.q or s.r ≠ k.r or s.l ≠ k.l      // k differs from s
11          S = S ∪ {s}; s = k                      // add s to S
12  S = S ∪ {s}                                     // add last s to S
13  return S                                        // S contains all MEMs
```

The above algorithm implements the extend-purge scheme for MEM computation. It is used for benchmarking the extend-purge approaches and is in accordance with previously published variants of this scheme for MEM computation [6-9]. The cited works mainly focus on strategies for cleverly selecting seeds for the set $K$. These selection strategies could be applied in the context of Alg. 1 (merge-extend based MEM construction) of the main manuscript as well. Hence, we do not incorporate or analyze these strategies here.

The algorithm consists of two segments:

Segment 1 (lines 1-5):

The for-loop visits all seeds in $K$ and maximally extends them to the left and right via the corresponding while-loops. This part of the algorithm is equal to the extension step in Alg. 1 (lines 12-16) of the main manuscript.

Segment 2 (line 6 -12):

This part deletes all duplicates resulting from the extension. First, we sort all seeds so that identical seeds become neighbors. Then we iterate over the sorted seeds and purge all seeds that are identical to their predecessor.

A comparison of the merge-extend and extend-purge strategies shows:

- The merge-extend benefits from long and "clean" reads, because the purge-extend strategy has to perform longer extensions for more minimizers ($k$-mers) here.
- The extend-purge strategy gains advantage over the merge-extend strategy in cases of few extensions because the purge step is computationally slightly less expensive than the merge step.

The above two observations are practically supported by the runtimes of Fig. 3 of the main manuscript as well as Supplementary Note 4.

### Supplementary Note 4 – Results for Illumina reads

#### a) Time Evaluation

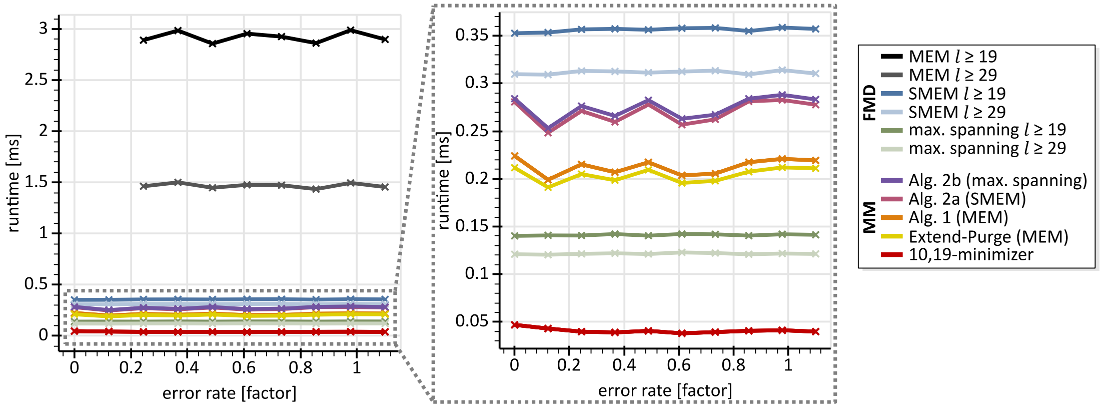


The diagram shows a runtime evaluation for Illumina reads similar to the runtime evaluation for PacBio CCS and CLR reads in Fig. 3 of the manuscript. Because the error rate of Illumina reads is quite low, the curves show a quite constant behavior. The curve for the extend-purge strategy is slightly below the curve of the merge-extend strategy, if the values are measured using the '-O2' optimization as shown in the above diagram. However, the order swaps if all compiler optimizations are switched off. Then, the curve for the extend-purge strategy is slightly above the curve of the merge-extend strategy. The curves are expected to be close to each other due to the short size of Illumina reads. Compared to Fig. 3B), the computation of maximal spanning seeds using the FMD-index is faster than all other variable-size seeding approaches. This is in accordance with the observation for maximal spanning seeds on error free reads in Fig. 3B).

#### b) Seed Entropy Analysis

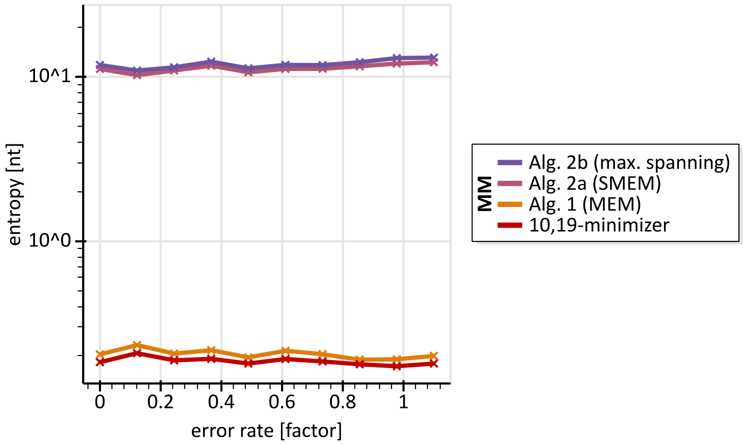


The diagram shows the seed entropy analysis for Illumina reads similar to the analysis for PacBio CCS reads in Fig. 4B). In accordance with the theoretical considerations with respect to the seed entropy, there is no change among the order of the curves for the respective kinds of seed sets (minimizer, MEM, SMEM, maximal spanning seeds). As for the runtime analysis, the curves express a quite constant behavior because the standard error for Illumina reads is quite low.

### Supplementary Note 5 – Comprehensive Occurrence Filter Effects

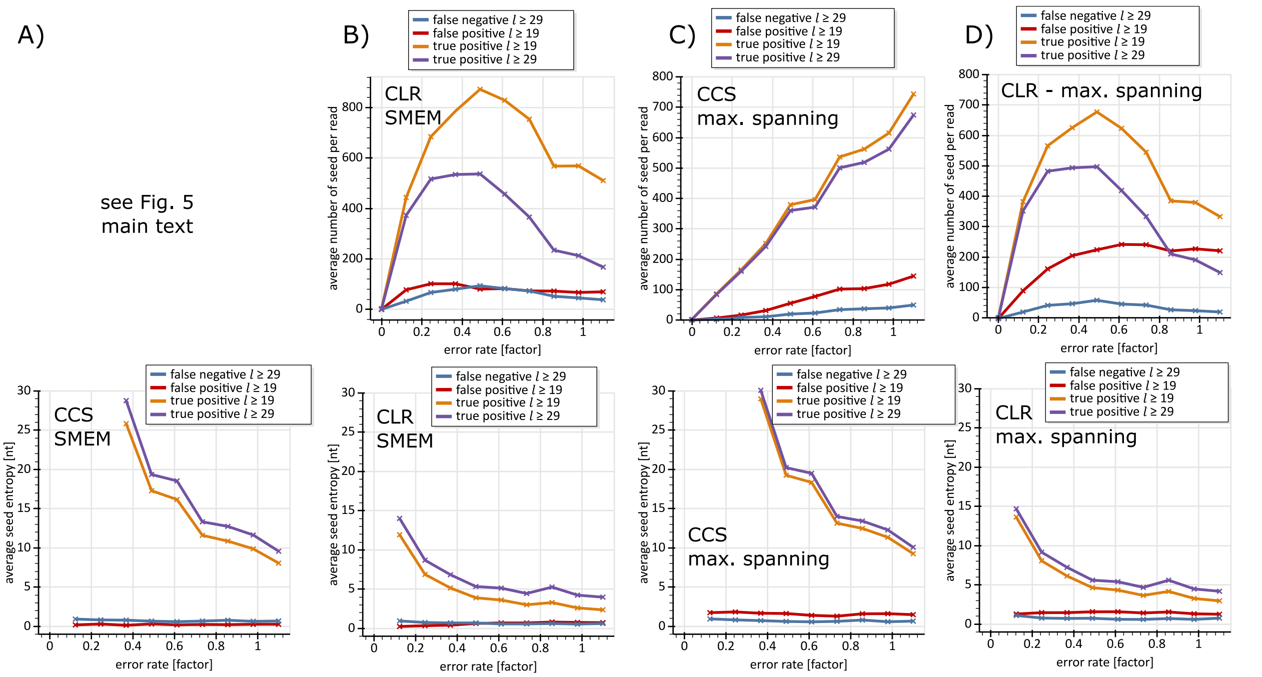


SMEM	Maximal spanning seeds

The above diagrams extend the occurrence filter analysis shown in Fig. 5 of the main manuscript: The left half of the eight diagrams show an analysis for SMEMs, while the right half show an analysis for maximal spanning seeds. The columns A) and C) show diagrams for CCS PacBio reads, while the columns B) and D) are for PacBio CLR reads. The y-axis in the top row expresses "number of seeds". Each point in the bottom row shows the average entropy of all seeds of its corresponding point (same color, same x-axis position) in the respective diagram in the top row. For low error rates (< 0.1), the entropy values are omitted for the following reasons:

- For true positives, the entropy tends towards the average read length.
- For false positives and false negatives, the lack of seeds turns the entropy meaningless.

The benchmarking environment and occurrence filter settings are described in the results section.

We first discuss the diagrams for CCS reads:

The curves for SMEMs and maximal spanning seeds are quite similar. Further, the number of false-positives (seeds erroneously identified as SMEMs by Alg. 2a or identified as maximal spanning seeds by Alg. 2b) and false-negatives (missed by Alg. 2a or Alg. 2b) are quite low. The entropy diagrams indicate that the false-negatives and false-positives are not relevant in the context of accurate alignments due to their low entropy.

We now discuss the CLR reads:

Because CLR reads have a worse quality than CCS reads, seeds are expected to be of shorter size than for CCS reads. With the decreasing size of seeds, the risk of a seed to occur multiple times on the reference increases. If the number of occurrences exceeds a given threshold (in the diagrams 200 occurrences), a seed is purged. This causes the decreasing behavior of the orange and purple curves in the top CLR diagrams, starting at an error rate of 0.5. Additionally, the entropy of the seeds close to an error rate of 1 is low for CLR reads compared with CCS reads. The number of false positives and false negatives for CLR reads does not differ significantly from the corresponding values for CCS reads with the exception of false positives in subfigure D). These false-positives need to be purged by the seed processing (chaining etc.).

#### Supplementary Note 6 – Runtime Evaluation

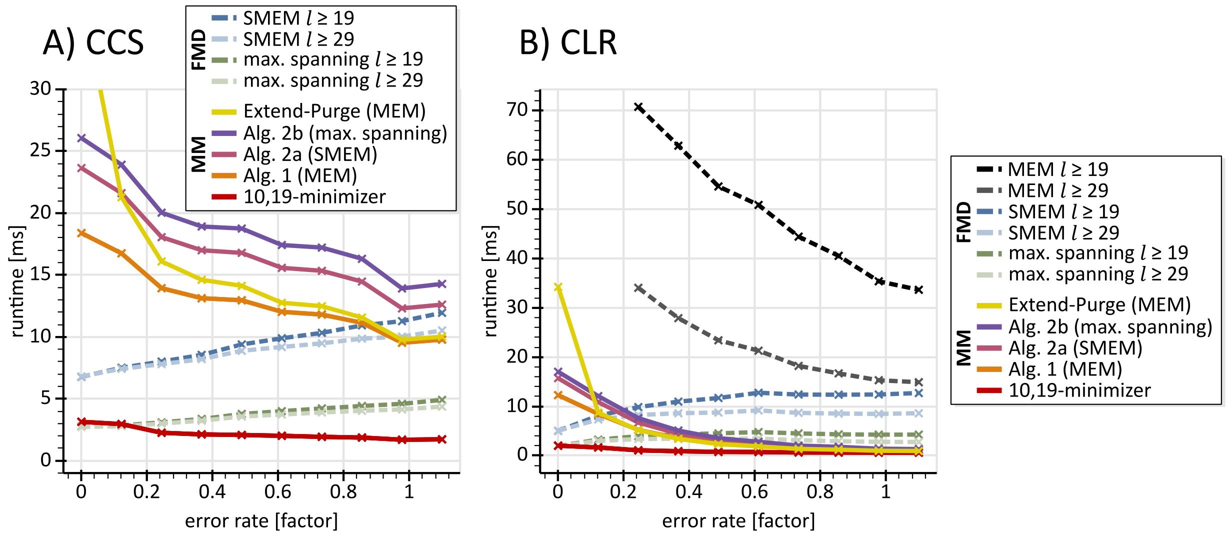


**A)** shows an extended form of diagram 3A) of the main manuscript. For CCS reads the SMEM and maximal spanning seed computation via the FMD-index is generally faster than via Alg. 2a or Alg. 2b. Please note that this superiority does not exist for MEMs.

**B)** shows an extended form of diagram 3B) of the main manuscript. As with CSS reads the generation of MEMs via the FMD-index is significantly slower than minimizer based approaches.
